# Giardia duodenalis: Base excision repair pathway enzyme FEN1 carries out catalytic activities pertaining to NER pathway

**DOI:** 10.1101/2025.04.23.650334

**Authors:** García-Lepe Ulises Omar, Tomás-Morales Sofía Gabriela, Izaguirre-Hernández María Teresa, Bazán-Tejeda María Luisa, Bermúdez-Cruz Rosa María

## Abstract

Giardia duodenalis is a binuclear protozoan that causes intestinal infection in humans and animals. The life cycle of G. duodenalis is comprised by 2 stages: trophozoite (vegetative, ploidy: 4N) and cyst (infective, ploidy: 8-16N) and the transition from one to another requires a precise coordination as well as the support of the DNA repair machinery. While DNA homologous recombination DNA repair has recently been characterized, NER and BER are pathways that had not been explored. Most of the structure specific enzymes (SSE) participate in a variety of processes like DNA replication stress, DNA adduct repair, Holliday junction processing. In an effort to explore these kinds of enzymes in *G.* duodenalis, a minimalist parasite, we aimed at characterizing the Fen1 enzyme by cloning its gene to study its catalytic properties (binding and nuclease) using flap and bubble DNA substrates. Unexpectedly, we found that GdFen-1 is able to cleave bubble DNA, then to shed light on which domains of this enzyme are responsible for this activity, giardial acid block and a portion of a cap region were substituted by their human counterparts, and while acid block substitution did not affect this activity, the modification in the cap region did. The possible implications of these findings are addressed.

**Highlights:**

- A functional homologue of the nuclease Fen1 is present in Giardia duodenalis
- Giardia duodenalis Fen1 nuclease binds and cleaves 5’ flaps
- Bubble-like structures are cleaved by the Fen1 protein of Giardia duodenalis

## 1. Introduction

Giardia duodenalis is an intestinal parasite that infects the upper small intestine causing acute watery diarrhea with 280 million cases in humans per year (Einarsson et al., 2015). This protozoan can infect all mammals and assemblages A and B have a zoonotic potential (transmitted from animals to human and vice versa). Giardial infection is present 2-3% in industrialized countries, while up 30% of its infections have been reported in developing countries. This parasite has two cycle stages: the trophozoite (vegetative stage) and the cyst (infective stage), with 4N and 8-16N ploidy, respectively, which is crucial for its survival (Bernander et al., 2001; Thompson et al., 1993). Thus, a precise coordination between stage transition and ploidy number are key for genomic stability and the DNA repair pathway is responsible for this task. Giardia has been shown to carry in its genome homologous recombination homologs and they have been shown to participate in DNA repair upon DNA damage infliction (Martinez-Miguel et al., 2017; Ordoñez-Quiroz et al., 2018; Sandoval-Cabrera et al., 2015; Torres-Huerta et al., 2016) however other DNA repair pathways such as the global genome repair (NER nucleotide excision repair) and BER (base excision repair) also very important during trophozoites growth have not been studied. The characterized structure specific endonucleases (SSE) (Dehé & Gaillard, 2017) pertaining to BER and NER pathways participate in various activities like DNA replication and transcription stress, DNA repair and Holliday structure processing. Among them, both members of the FEN1–XPG family are able to cleave flap DNA, however only XPG is able to cleave bubble DNA. Since *Giardia duodenalis* pathways are atypical and/or minimalist (Morrison et al., 2007), we undertook the study of GdFen1. In this work we aimed at studying Fen1 (Flap endonuclease 1) by cloning the gene, expressing and purifying the protein to analyze its binding activity using DNA substrates either with flap or bubble structures and then its nuclease activity using these substrates. Interestingly, we found that GdFen-1 is able to cleave both flap and bubble DNA structures. Then, to identify domains responsible for this bubble cleavage activity, we substituted either the giardial acid block or a portion of the giardial cap region by its human counterparts and found that while human acid block modification in GdFen1-HsAB did not affect this activity, the human cap domain modified did. We discuss the possible reasons as well as the implications for these activities.

## 2. Materials and methods

### 2.1 Cell culture

Trophozoites of Giardia duodenalis (Assemblage A1, WB strain, ATCC 30957) were grown at 37° C in 50 mL conical tubes in modified TYI-S-33 medium (Keister, 1983), supplemented with 10% fetal bovine serum (HyClone) and 1X antibiotic/antimycotic solution (Hyclone, Thermo Scientific SV30079.01).

### 2.2 Bioinformatics analysis

The nucleotide sequence of GdFen1 was obtained from GiardiaDB (https://giardiadb.org/giardiadb/app). GdFen1 ID: GL50803_16953. Additional sequences from Fornicata organisms used to perform multiple sequence alignment (MSA) were obtained using BLAST (NCBI)(Sayers et al., 2024). Multiple sequence alignment (MSA) was made using MAFFT software (Katoh et al., 2013) and visualized through the ESPript 3.0 program (Gouet et al., 2003). To identify functional domains, GdFen1 protein sequence was scanned in the NCBI conserved domains software (https://www.ncbi.nlm.nih.gov/Structure/cdd/wrpsb.cgi) (Marchler-Bauer et al., 2015). The 3D structural organization was predicted through AlphaFold2 software (Jumper et al., 2021) and subsequently aligned and visualized with their human homologue through UCSF ChimeraX (Meng et al., 2023; Pettersen et al., 2021). The TM-score and RMSD were calculated through the online software TM-align (Zhang & Skolnick, 2005)

### 2.3 Trophozoites genetic material extraction, gene amplification and cloning

Trophozoites from a confluent culture were collected by centrifugation, washed with PBS 1X and lysed with lysis solution (10 mM Tris-HCl pH 7.4, 10 mM EDTA, 150 mM NaCl, 0.4% SDS and 20 mg/mL proteinase K) for 16 hours at 65° C, subsequently, the mix was treated with 20 µg/mL RNAse A enzyme for 1 hour at 37° C. The extraction was made with 1:1 phenol-chloroform-isoamyl acid and precipitated with 2.5 volumes of absolute ethanol and 10 % of 3 M sodium acetate. An endpoint PCR reaction was performed to amplify GdFen1 directly from genomic DNA (this gene does not possess intronic sequences). To facilitate cloning, an intermediary commercial vector pJet 1.2/blunt (Thermo Fisher Scientific K1231) was used and specific restriction sites were added to the 5’ end of primers to facilitate cloning (Table 1) GdFen1 containing fragment was cleaved out with *Sac*I and *Nhe*I restriction enzymes. The obtained DNA fragment was purified and cloned into the inducible expression plasmid pET- 100/D-TOPO (Invitrogen K10001) using T4 DNA ligase (NEB Inc. M0202), generating the pET100-GdFen1 plasmid (Figure 2b). The plasmid includes a polyhistidine tag at the amino end of the recombinant protein, employed to detect and later purify the protein. Plasmid map was built using the software Vector NTI 11.5 (Invitrogen).

**Table 1.**
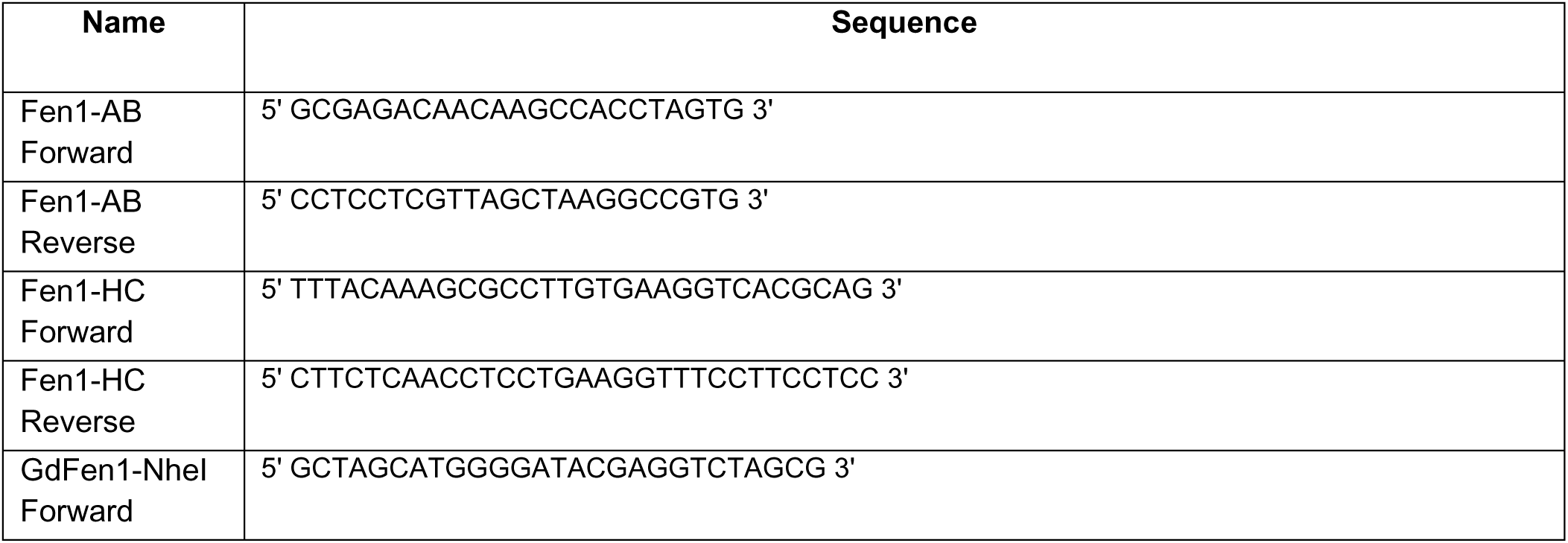
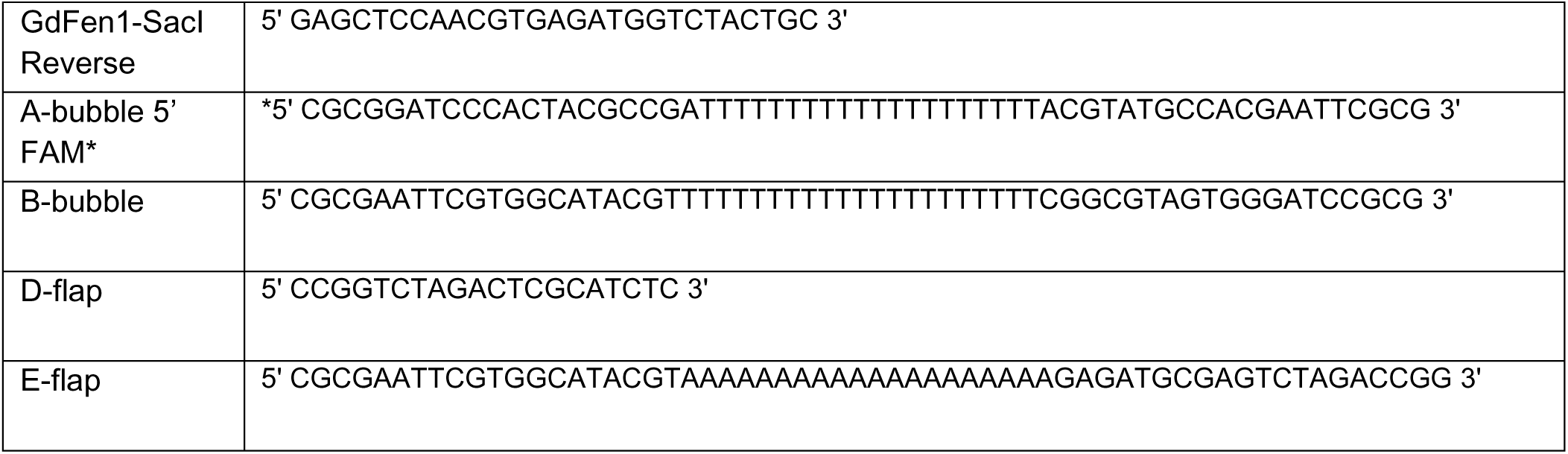
Oligonucleotides used in this study.

### 2.4 Generation of mutants

Two GdFen1 mutants were created from the pET100-GdFen1 plasmid using the Q5^®^ Site- Directed Mutagenesis Kit (NEB E0554S). The acid block or a portion of the helical cap sequences were substituted with corresponding human counterparts to generate pET100- GdFen1-HsAB and pET100-GdFen1-HsHC plasmids respectively. The pET100-GdFen1 plasmid was amplified through PCR using Q5 Hot Start polymerase and specifically designed oligonucleotides bearing the desired sequence change and 5’ ends annealing back-to-back (see Table 1), after amplification the plasmids were incubated with KLD Enzyme Mix and next transformed into Escherichia coli XL1-Blue competent cells. The correct incorporation of the mutation was corroborated by DNA sequencing.

### 2.5 Protein expression and purification

The plasmids were transformed into the SoluBL-21 E. coli strain (amsbio C700200). One isolated colony was grown in LB media added with 100 μg/ml of ampicillin and incubated overnight at 37° C. This culture was used to inoculate a second one which was induced with 1 mM isopropyl β-D- 1-thiogalactopyranoside (IPTG) and incubated for up to 6 h at 37°C. The culture was cooled, and the bacterial pellet was obtained by centrifugation. Pellets were suspended in lysis buffer (7.53 mM Tris-base pH, 2 M NaCl, 20 mM MgCl2), sonicated (12 cycles of 5 seg at 100% with intervals of 35 seg) and centrifuged (14,000 RPM for 35 min). The supernatant was recovered and settled into an affinity chromatography column with HisPur Ni-NTA superflow agarose beds (Thermo Fisher Scientific 25 215), washed with lysis buffer with low imidazole concentration (50 mM), and eluted with a 300 mM imidazole solution. The eluted solution was concentrated using an Amicon Ultra-4 Column (Millipore UFC803024), and quantified by the colorimetric method of Bradford, using the Bio-Rad Protein Assay Dye Reagent Concentrate (BIO-RAD 500-0006). Solid-phase immunodetection was performed with 1:1000 primary anti-polyhistidine antibody (Sigma-Aldrich H1029), and 1:15000 secondary anti-mouse IgG (H + L) HRP conjugate (Promega W4021).

### 2.6 Probe hybridization

Two different fluorescent probes were designed: Bubble probe, equivalent amounts of primers: A-bubble 5’ FAM and B-bubble were mixed, incubated at 80° C for 5 min and allowed to cool at room temperature for 2 h. Both primers contain the same polythymine sequence in the middle to avoid hybridization and thus form a bubble structure. Flap probe, equivalent amounts of primers: D-flap and E-flap were incubated as mentioned before, then, a fluorescent-tagged primer A (A-bubble 5’ FAM) was added to the mixture and incubated at 50° C for 5 min, then it was allowed to cool at room temperature. Both probes, Bubble and Flap, harbor a FAM (6- carboxyfluorescein) molecule attached to the 5’ end. The correct hybridization pattern was corroborated through polyacrylamide gel electrophoresis (data not shown). See Table 1 for sequences used to amplify the GdFen1 gene and to generate the probes.

### 2.7 DNA binding assay

Increasing concentrations of purified GdFen1 were incubated with Flap or Bubble fluorescent probe in binding buffer (40 mM Tris-HCl, 1 mM DTT 1, 5 mM MgCl2). The reactions were loaded and resolved in a 4 % native polyacrylamide gel in TAE buffer (40 mM Tris-acetate, pH 7.5 and 0.5 mM ethylenediaminetetraacetic acid). The probes and their complexes with GdFen1 were visualized in an Odyssey FC Imaging System (LI-COR Biosciences) at 600 nm. The observed bands were measured by densitometry utilizing the Image Studio Lite Version 5.2 software (LI-COR Biosciences).

2.8 DNA nuclease assay

To determine nuclease activity, a fixed concentration of purified protein GdFen1 was incubated with the Flap or Bubble fluorescent probe in nuclease buffer (30 mM Tris-HCl, 1 mM DTT, 50 mM KCl, 50 μg/ml BSA, 5mM MnCl2) for 30-, 60-, 90-, 120-, or 180-min as indicated. After incubation, the reactions were stopped with 0.1 mg/ml proteinase K and 3% SDS and resolved in a 10 % native polyacrylamide gel in TAE buffer. All of the electrophoresis were run under the same conditions of time and voltage, 140 V for 40 min. The nuclease activity detection, visualization and densitometry measurement were conducted as was previously mentioned.

## 3. Results

### 3.1. Conserved domains in GdFen1

The nucleotide sequence of the Giardia duodenalis Fen1 (GdFen1) was obtained from the GiardiaDB informatics resources and translated to perform the following in silico analysis. Multiple sequence alignments (MSA) were employed to examine the degree of conservation and the presence of functional domains. The giardial protein sequence was aligned with homologues from various mammal models and other closely related organisms, including some parasites in the Fornicata phylum. Through the use of pairwise sequence analysis, it was determined that GdFen1 shares an amino acid sequence identity of 40.2% with its human counterpart. Figure 1a shows a simplified MSA of GdFen1, highlighting the functional domains relevant to protein activities, which were identified using the NCBI conserved domains software (Marchler-Bauer et al., 2015), as well as other secondary structure elements relevant for protein function. Additionally, the secondary structure information of GdFen1 was also explored using the crystallized protein structure of human Fen1 as a template (HsFen1, PDB 3q8k) (Tsutakawa et al., 2011)and included in upper part of the MSA. As can be observed, despite the modest sequence identity between human and Giardia sequences, the XPGN, XPGI and DNA binding domains are well conserved, suggesting that GdFen1 may present the activities associated with the Fen1 nuclease in other organisms. The black square in Figure 1a denotes “the acid block” (solid lines), a small glutamic acid-rich sequence that has been associated with substrate preference among structure-specific endonucleases (SSE) (Tsutakawa et al., 2011). Interestingly, this small sequence is not completely conserved among Fornicata, since half of the glutamic acid residues have been replaced by other non-acidic residues. On the other hand, the black square with dotted lines indicates a segment of the “the helical cap”, a sequence suggested to be implicated in nuclease preference for the 5’ ends and formation of the helical gateway (Tsutakawa et al., 2011). The acid block, the helical cap and their implications in the Giardia’s Fen1 are discussed in more detail below. A global streamlined MSA is depicted in Figure 1b, illustrating that GdFen1 shares an important degree of conservation with closed orthologs in terms of the presence of the amino acid domains and overall length. As can be noted the carboxyl terminus is the part of the protein that shows the most variability among the orthologs included in the analysis, but no relevant motifs or domains were found using the NCBI tool or described elsewhere. The tertiary structure analysis of GdFen1 reveals that GdFen1 also shares the same overall fold with the human FEN1, with a TM-score of 0.87 and RMSD of 1.9, as can be noted in Figure 1c. These in silico results strongly suggest that the G. duodenalis parasite retains a FEN1 homologue that could have the functions described in other organisms, due to its high degree of conservation in functional domains as well as its three-dimensional folding, which is very similar to its human homologue.

**Figure 1.**
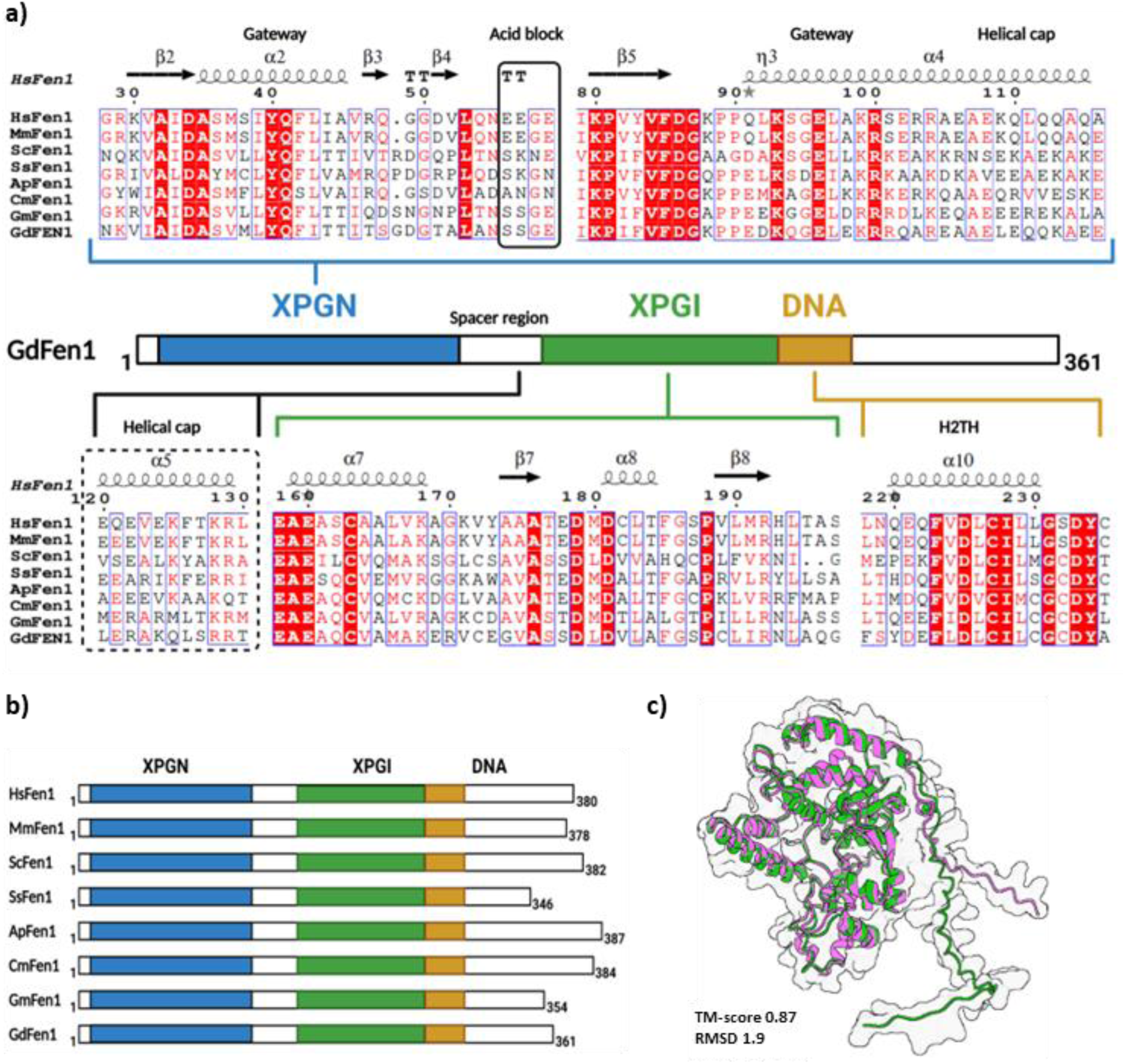
In silico analysis of the GdFen1 protein sequence. a) A simplified multiple sequence alignment (MSA) of GdFen1 with other model and related organisms was generated, and the major predicted functional domains are shown. The acid block and a segment of the helical cap are framed in solid and dotted squares respectively. Hs, Homo sapiens (NP_004102.1); Mm, Mus musculus (P39749.1); Sc, Saccharomyces cerevisiae (NP_012809.1); Ss, Spironucleus salmonicida (KAH0577767.1); Ap, Aduncisulcus paluster (GKT28123.1); Cm, Carpediemonas membranifera (KAG9391069.1); Gm, Giardia muris (TNJ28849.1); Gd, Giardia duodenalis (GL50803_16953). b) Global MSA showing the functional domains and the total length of the amino acid sequences. c) 3D alignment of the predicted tertiary structures of GdFen1 (shown in pink) with the human homologue (shown in green). The TM-score and RMSD are indicated.

### 3.2 Gene cloning and recombinant protein purification

To confirm our bioinformatic predictions and to characterize the functions of GdFen1, the corresponding gene was cloned, and the recombinant protein was expressed and purified using an E. coli bacterial system. Despite being a eukaryote, only a few G. duodenalis genes are known to contain introns (Morrison et al., 2007), therefore the GdFen1 gene was amplified directly from G. duodenalis genomic DNA using endpoint PCR, without requiring the cDNA generation step in this case. Consistent with our in silico analysis, the resulting amplicon was 1,112-bp long, as shown in Figure 2a. The GdFen1 gene was first subcloned into the pJet 1.2/blunt plasmid (see Materials and Methods for details) and later introduced into the pET-100 expression vector to create the pET-100-GdFen1 plasmid (Figure 2b). This vector has a T7 RNA polymerase promoter and contains a polyhistidine tag fused to the N-terminus of the protein, the T7 RNA polymerase expression is IPTG-induced in the BL21(D3) strain used. This system allows verification of protein expression by solid-phase immunodetection, as shown by the kinetic expression of GdFen1 (Figure 2C), where an anti-His antibody was used to detect the predicted 41 kDa protein. During the induction process with 1 mM IPTG, the expression of GdFEN1 expression was observed to increase with the incubation time, as shown by the higher amounts of the protein in the samples (Figure 2c). Additionally, the tag facilitates protein purification via affinity chromatography, as can be observed in the Coomassie-stained polyacrylamide gel where the purified protein is compared with a total bacterial extract (TE) prior to purification in lane 2 (Figure 2d). These data confirm that the G. duodenalis genome bears the Fen1 gene that was predicted by bioinformatics, and that the gene is functional since it produces a protein with the predicted molecular weight.

**Figure 2.**
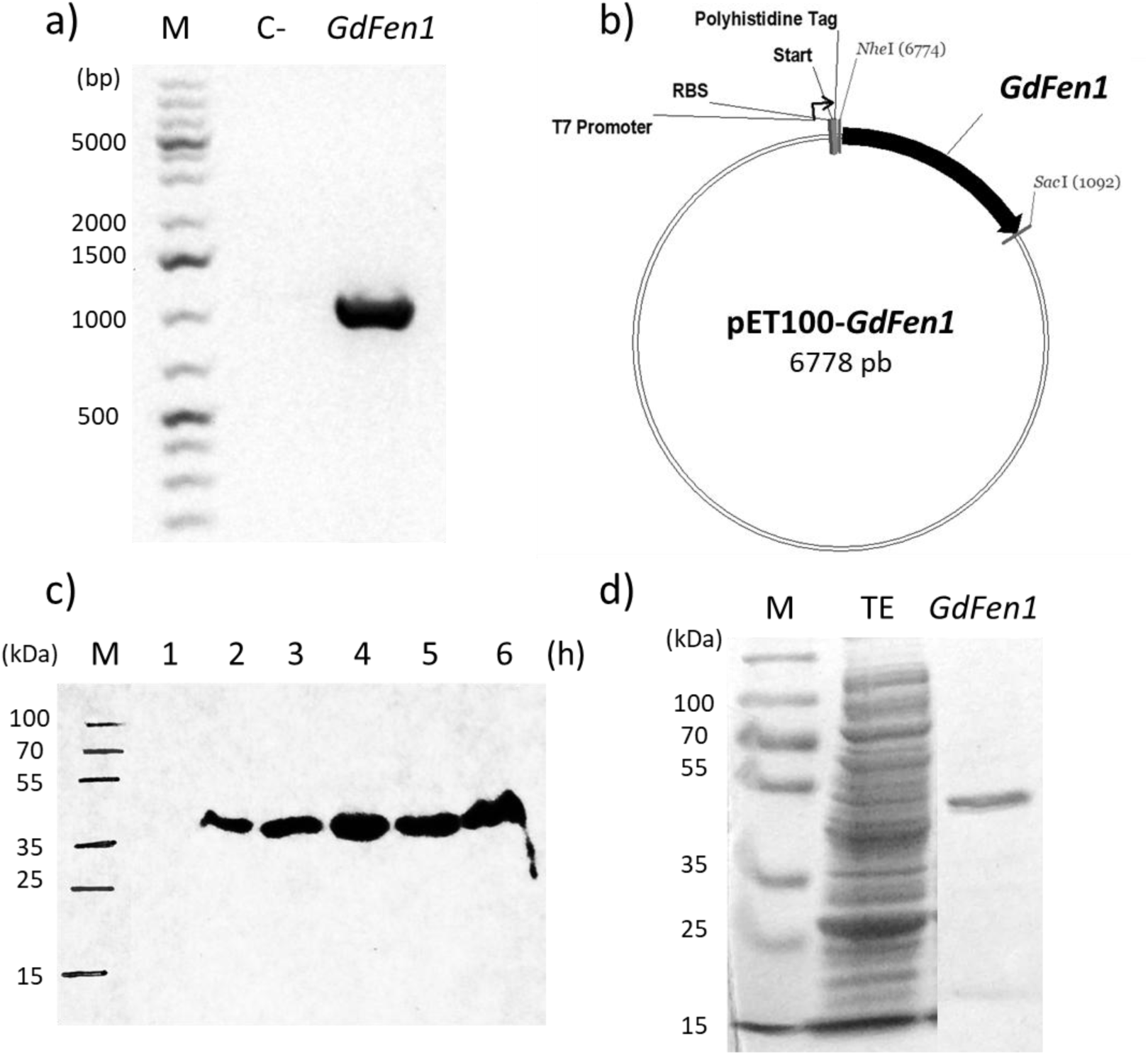
Gene cloning and protein expression. a) 1% agarose gel showing the endpoint PCR amplicon of the GdFen1 gene (1112-pb) amplified from G. duodenalis genomic DNA, along with a negative control (C-). b) An image of the expression plasmid for GdFen1, pET100- GdFen1. c) An image showing the solid phase immunodetection of GdFen1 expression kinetics (41 kDa) using an anti-His antibody. d) A 12% polyacrylamide gel electrophoresis composite image stained with Coomassie dye, showing total bacterial extract (TE) in lane 2 and the purified GdFen1 in lane 3 (the other lanes of the gel, including the washes and the first purified fractions, have been cut out). In images a), c) and d) molecular weight markers (M) are shown at the left.

### 3.3 GdFen1 interacts with flap structures and exhibits endonuclease activity

Fen1 nuclease is involved in DNA repair and replication processes, where it uses this activity to cleave off 5’ ends in a sequence-independent manner but structure-dependent, allowing the repair machinery to continue the process and seal the DNA double strand (Lieber, 1997; Liu et al., 2004). In this regard, to first evaluate whether the *Giardia* homologue conserves the ability to bind DNA flap structures, the purified recombinant GdFen1 was tested through electrophoretic mobility shift assay (EMSA). Increasing concentrations of protein were incubated with a double-stranded oligonucleotide (60-bp) bearing a flap structure of 20-bp (from now on the Flap probe) labeled at the 5’ end with the fluorescent tag 6-carboxyfluorescein (Figure 3a, top section). Protein-DNA interaction is visualized as a shift of the probe fluorescent band (free probe). This essay confirmed the ability of this protein to bind the specific flap DNA structure, as shown in Figure 4a. The fluorescent bands representing DNA-protein complexes were quantified by densitometric analysis and plotted (Figure 4b). In the same way, the ability of GdFen1 to cleave the flap structure through endonuclease activity was evaluated. The purified recombinant protein was incubated with the Flap probe in nuclease buffer (see Materials and Methods) for increasing time points. During the reaction, the 5’-end of the Flap probe is cleaved by GdFen1, releasing a fragment 20-bp long containing the fluorescent tag (Figure 3a, lower section). This smaller product is observed at the bottom of the polyacrylamide gel, while the intact probe at the top decreases with increasing incubation time (Figure 4c). The residual (non-processed) probe was quantified and analyzed by densitometry (Figure 4d). The above results clearly demonstrate the ability of the Giardial Fen1 homologue to process 5’ flap structures, suggesting that this minimalist organism relies on the repair pathways necessary to process flap intermediates during replication and/or DNA repair.

**Figure 3.**
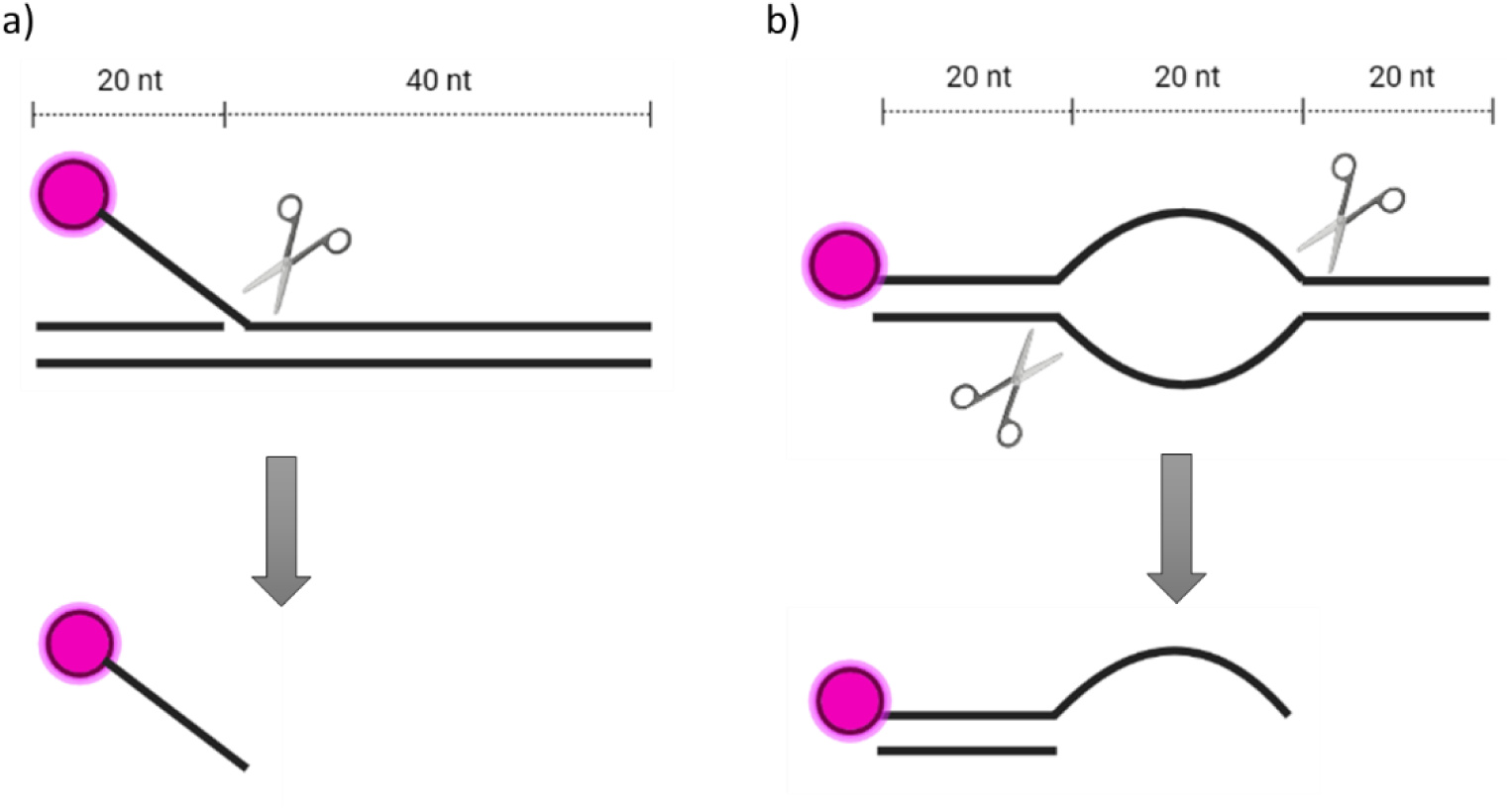
Schematic of the fluorescent probes used in this study. a) Top, Flap probe contains a 20-nt flap end labeled with the fluorescent tag 6-carboxyfluorescein at its 5’ end. Bottom, the fluorescent product obtained cleaving off the flap. b) Top, Bubble probe contains a 20-nt bubble structure in which only one of the two oligonucleotides used is labeled at its 5’ end with the fluorescent tag. Bottom, the fluorescent product obtained after processing the bubble by cutting them at the 3’ ends. Created with BioRender.com

**Figure 4.**
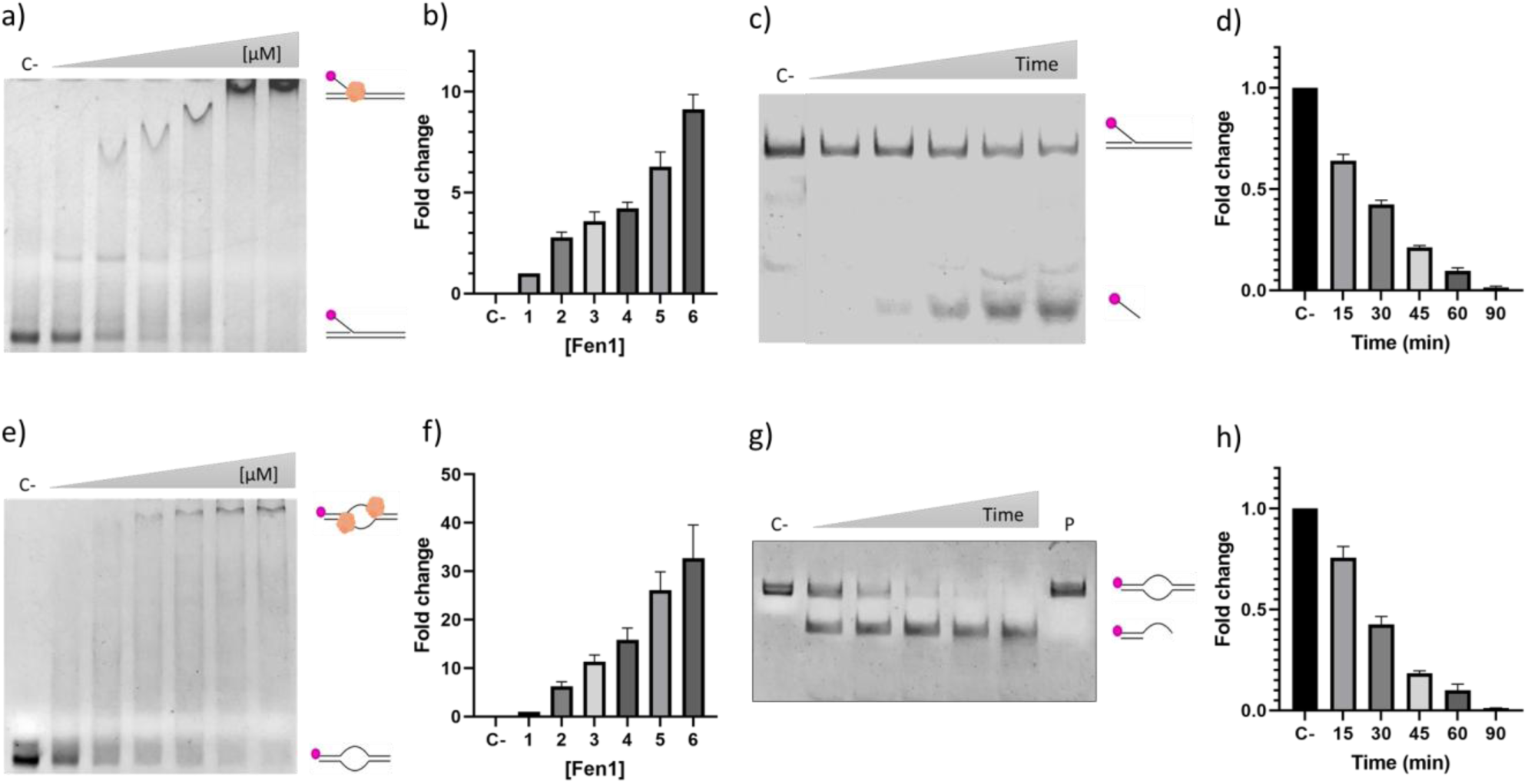
Binding and cleavage of the Flap and Bubble probes by GdFen1. a) Fluorescence EMSA with increasing concentrations of purified GdFen1, shifted DNA-protein complexes and free probes are shown. b) Densitometric analysis plot of the DNA-protein complexes from a) normalized against the lowest protein concentration. c) Time course of nuclease activity of GdFen1 on Flap probe. d) Densitometric analysis plot normalized against the negative control (no protein) from c). e) Fluorescence EMSA using increasing concentrations of purified GdFen1, shifted DNA-protein complexes and free probes are shown. f) Densitometric analysis plot of the DNA-protein complexes from d) against the lowest protein concentration. g) Time course of nuclease activity of GdFen1 on the Bubble probe. h) Plot of the densitometric analysis normalized with the negative control (no protein) from g). Error bars represent SD.

### 3.4 GdFen1 is able to bind and cleave DNA bubble structures

Fen1 is a member of the Fen1 family of proteins (Dehe & Gaillard, 2017). This group includes other homologous nucleases, such as the XPG, a significantly larger protein that conserves the same two fundamental domains (XPGN and XPGI) involved in the nuclease activity (Fagbemi et al., 2011). In contrast to Fen1, XPG participates in the nucleotide excision repair (NER) pathway, which typically acts on bulky lesions that alter the spatial conformation of the DNA, such as DNA-adducts or typically those produced by UV light (Spivak, 2015). Notably, although the use of UV irradiation on trophozoites and cysts is an accepted method to eradicate them with several purposes (Adeyemo et al., 2019; Li et al., 2008; Linden et al., 2002), the presence of the NER pathway has not been characterized in G. duodenalis, and indeed it has been suggested that G. duodenalis does not possess the protein homologues necessary to perform this task (Einarsson et al., 2015). Interestingly, some potential homologues of NER proteins have been found by our research group using bioinformatic approaches (data not shown), XPG among them. Because of the conservation of XPGN and XPGI domains in GdFen1 and that Giardia duodenalis went through a reduction of complexity due to adoption of a parasitic lifestyle (Cernikova et al., 2018), we asked whether GdFen1 has the ability to bind and cleave bubble DNA (generally processed by NER pathway enzymes). To evaluate this hypothesis, an EMSA was performed with a Bubble probe, a specific substrate for the XPG nuclease, another member of the Fen1 family (Scharer, 2013), which contains a 20-nt bubble in the middle zone of the probe, consisting of only T nucleotides in both strands to avoid base pairing, and a fluorescent tag at one of its 5’ ends (Figure 3b, top section). As can be observed, GdFen1 is able to bind the Bubble probe. This is demonstrated by the appearance of a band corresponding to protein-DNA complexes at the top of the gel as the protein concentration increases (Figure 4e). The fluorescent bands representing DNA-protein complexes were quantified by densitometric analysis and plotted (Figure 4f). To further investigate this observation, a nuclease assay was performed using the Bubble probe. This probe is designed so that one of its products retains the fluorescent label upon cleavage at the 3’ end of the bubble (since this is the end where XPG cleaves), so that processing can be detected by a change in the size of the probe (Figure 3b, lower section). Unexpectedly, it was observed that GdFen1 is able to cleave bubble structures, as can be seen in Figure 4g with the formation of a shorter product as time increases. The residual (non-processed) probe was quantified and analyzed by densitometry (Figure 4h). The products generated when the Bubble probe is cleaved or processed are evidently larger than those generated from the Flap probe, thus confirming that the bubble structure is being twice cut in both 3’ ends, since a single cut would not produce probe fragmentation due to the base pairing on the edges of the probe (Figure 4c and g). These observations additionally confirm that the two probes employed in this investigation have the expected structures and are cleaved as designed (Figure 3a and b). Notably, as the cutting of the Bubble probe was a peculiar observation, the experiment was repeated with another bubble probe generated with a completely different set of oligonucleotides, where GdFen1 was also able to cleave (data not shown). In summary, these observations demonstrate that GdFen1 is able to bind and cleave bubble structures in vitro, features commonly associated with the XPG protein of the NER pathway, raising the possibility that GdFen1 may be also involved in the unexplored NER repair pathway in G. duodenalis.

### 3.5 The helical cap is involved in GdFen1 bubble cleaving activity

As was demonstrated in the previous results, GdFen1 not only shows affinity and processing on flap-like substrates, but notably it is also able to bind and cleave bubble-like structures, which are normally processed by XPG of the NER pathway. Both proteins have the same cleavage domains, but one of the major differences between them is the presence of a so-called Spacer region or R-domain sequence in XPG, which is presumed to help separate the functional domains so that they can process larger structures such as bubbles (Hohl et al., 2007; Tsutakawa et al., 2011). Clearly, this sequence of about 680 amino acids in the human protein is not present in GdFen1 or other Fen1 in different organisms (Figure 1b, c), thus there must be some other feature that allows Fen1 to process this type of structure. Comparison of GdFen1 with its human counterpart revealed that the acid block and the second segment of the helical cap are not completely conserved in the parasite homologue (Figure 1a). Besides, the second segment of giardial helical cap (α5 helix) exhibits a different charge distribution since it bears more positively charged amino acids and less negatively charged than the human protein (Figure 1a). Interestingly, this segment of the helical cap is not conserved in the human XPG (Tsutakawa et al., 2011), which may indicate this region is not fundamental for bubble cleavage. The acid block and the helical cap in HsFen1 have been directly linked to the substrate specificity of the enzyme, the first one acting as constraint limiting the cleavage of other structures, and the helical cap as a fundamental structural element for the formation of the helical gateway, the entrance to the main cavity where the catalytic activity takes place (Tsutakawa et al., 2011). To confirm whether the differences of these sequences in GdFen1 are responsible for its unexpected behaviour, the acid block and helical cap sequences (Figure 1a) were separately replaced by their human equivalents by site-directed mutagenesis in the pET100-GdFen1 plasmid, generating the pET100-GdFen1-HsAB and pET100-GdFen1-HsHC plasmid mutants. The corresponding proteins GdFen1-HsAB and GdFen1-HsHC carry the human acid block and helical cap, respectively. Both GdFen1 mutants were expressed, and their biochemical properties were evaluated as previously done with wild-type GdFen1. Neither mutation affects the ability to bind to the bubble probe, as seen in the EMSA assays where the retarded bands, indicating interaction with the DNA, are detected at the top of the gel (Figure 5a and c). For further investigation, the nuclease activity of the GdFen1 mutants was assessed using either the Bubble or Flap probes. The exchange of the acid block does not alter the nuclease ability of GdFen1 over the bubble or flap probes, suggesting that this small segment of the protein is not directly involved in substrate determination in the giardial protein (Figure 5e and g, AB lanes). On the other hand, when the helical cap was exchanged with its human counterpart, the nuclease activity on the Bubble or Flap probes was almost completely abolished (Figure 5e and g). These observations strongly suggest that this segment of the helical cap is directly involved in the ability of GdFen1 to process bubble structures, while the acid block was irrelevant for cleaving.

**Figure 5.**
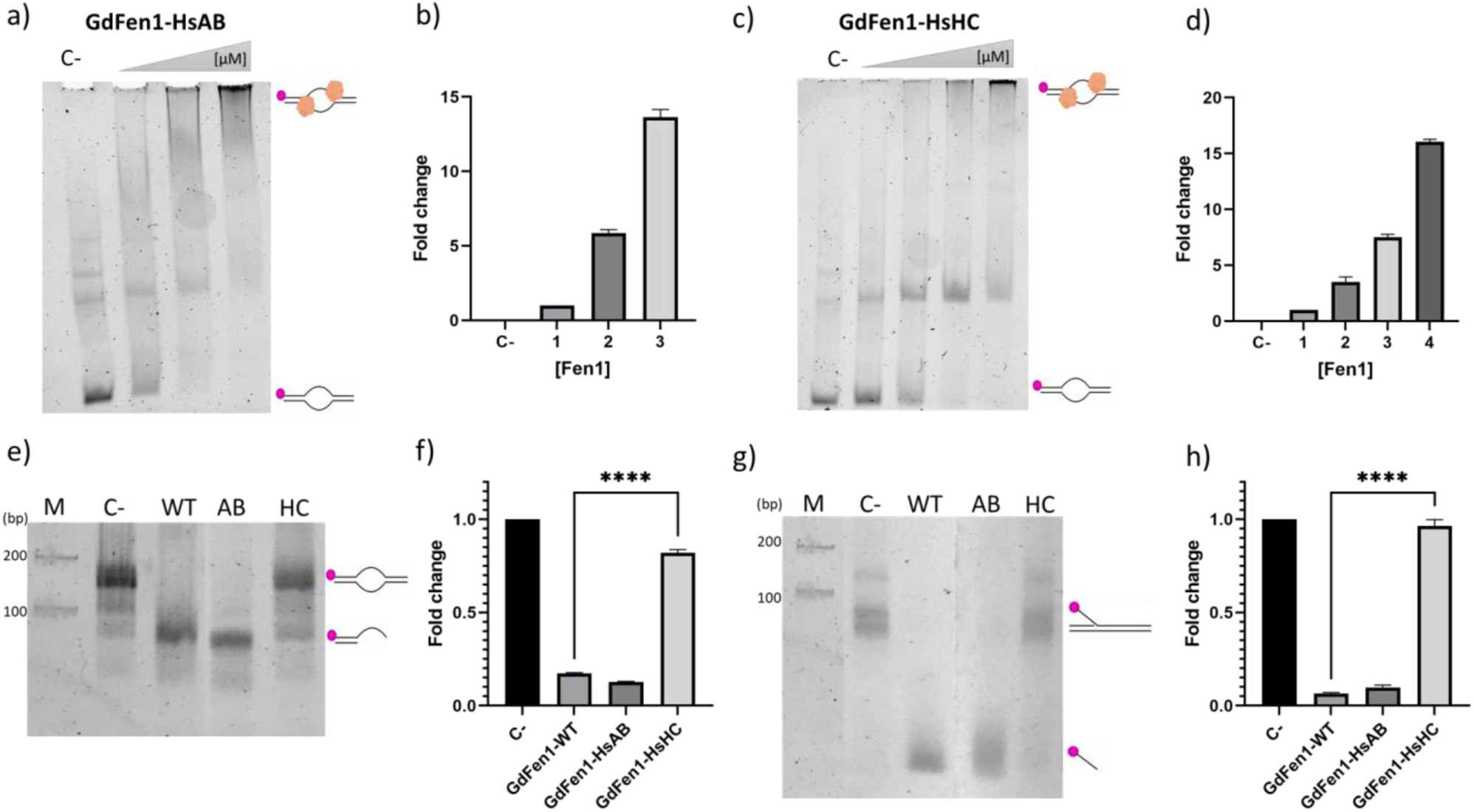
Binding and cleavage of the Bubble and Flap probes by GdFen1 mutants. a) Fluorescence EMSA with increasing concentrations of purified GdFen1-HsAB, shifted DNA-protein complexes and free probes are depicted. b) Densitometric analysis plot of the DNA-protein complexes from a) normalized against the lowest protein concentration. c) EMSA with the mutant GdFen1-HsHC, shifted DNA-protein complexes and free probes are shown. d) Densitometric analysis plot of the DNA-protein complexes from c) normalized against the lowest protein concentration. e) Nuclease activity over the Bubble probe: M, molecular marker; C-, no protein control; AB, acid block mutant (GdFen1-HsAB); HC, helical cap mutant (GdFen-HsHC). f) Densitometric analysis plot of the residual (non-processed) probe, normalized against the negative control (no protein) from e). g) Nuclease activity over the Flap probe. h) Densitometric analysis plot of the residual (non-processed) probe, normalized against the negative control (no protein) from g). Error bars represent SD. Statistical analysis was performed using a one-way ANOVA test. ****P < 0.0001 compared to GdFen1-WT, as indicated by the black lines.

### 3.6 The helical cap of GdFen1 enlarges the gateway cavity

With the intention of exploring the previous observations in more detail and determining whether these mutations affect the tertiary structure of the protein in any way, the structure of the mutant proteins was predicted using AlphaFold2 (Jumper et al., 2021). Figure 6 shows the structures of HsFen1, GdFen1, GdFen1-HsAB and GdFen1-HsHC, with the acid block and helical cap highlighted in black and red, respectively. As these mutations only humanise the giardial protein in some way, no major changes in the overall tertiary structure are observed, however, when a cavity determination analysis was performed (Guerra et al., 2021), it was found that in wild-type GdFen1 the main cavity is enlarged compared to HsFen1 (Figure 6d and e). Notably, when the helical cap is mutated, the volume of the cavity is significantly reduced from 637.63 Å^3^ to 572.18 Å3 and takes on a shape and volume (511.92 Å^3^) similar to the human protein. This effect could explain why the GdFen-HsHc mutant loses the ability to process bubble structures when the segment of the helical cap is exchanged for the human version (Figure 5e). On the other hand, the mutation in GdFen-HsAB, which modifies the giardial acid block, does not induce a significant change in the conformation or volume of the cavity, consistent with the conservation of the ability to process bubble structures (Figure 5e).

**Figure 6.**
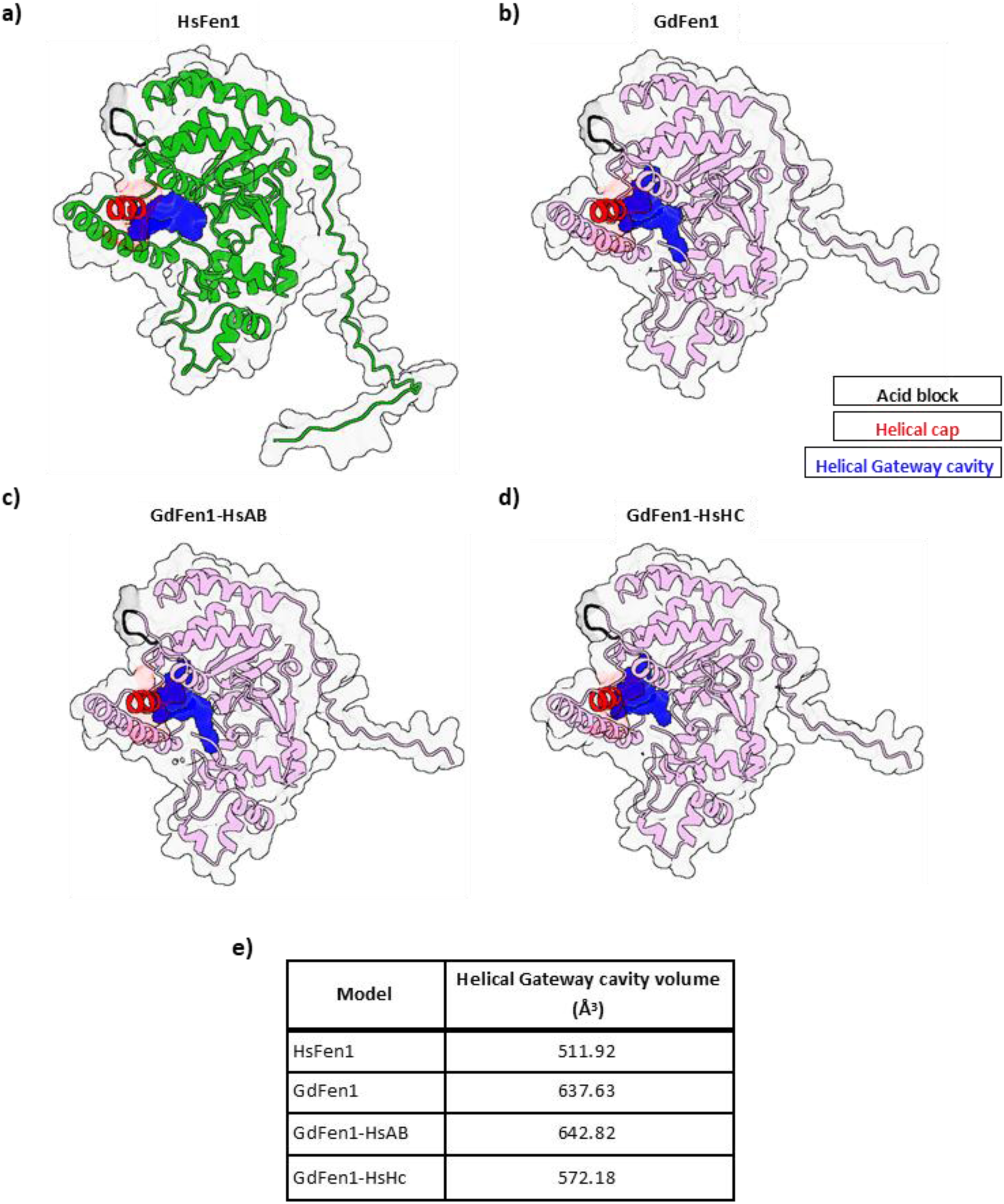
Structure determination and analysis of Fen1 orthologs and mutants. All the models depicted were generated through AlphaFold2. The acid block and helical cap are highlighted in black and red, respectively. The helical gateway cavity is shown in blue. a) HsFen1. b) GdFen1. c) GdFen1-HsAB. d) GdFen1-HsHC. e) Table with the cavity analysis results, the volumes of the main cavity are shown in Å^3^.

To complement our study and explore the implications of the helical cap on the distribution of charges on the protein surface, we determine the Coulombic Electrostatic Potential (ESP) of HsFen1, GdFen1 and their mutants using the ChimeraX built-in analysis tool. The results are shown in Figure 7. As can be seen, GdFen1 has a more positive surface area in the DNA interaction region compared to the human protein (Figure 7a and b, dotted rectangles). Similarly, the region containing the fragment of the helical cap (α5 helix) shows a more positive surface. In HsFen1-HsAB, as expected, the mutation enhances the negativity of the acid block, but the ESP remains unchanged in the rest of the structure (Figure 7c) dotted circle). Interestingly, the inclusion of the human helical cap in GdFen1 generates a stronger negative surface around the helical gateway. This observation may explain why the GdFen1 HsHC mutant loses nuclease activity, as the helical gateway and thus the active site may be inaccessible to DNA due to charge repulsion. Furthermore, the electrostatic surface on the rest of the protein is not affected, which explains the conservation of DNA binding activity (Figure 7d dotted circle). Taken together, these findings suggest that the unique features of the GdFen1 helical cap influence the volume and conformation of the helical gateway cavity, allowing it to process not only flap but also bubble structures. In addition, the overall sequence of GdFen1 creates a more positive surface at the DNA-interacting region, which may contribute to the atypical catalytic activities of the protein.

**Figure 7.**
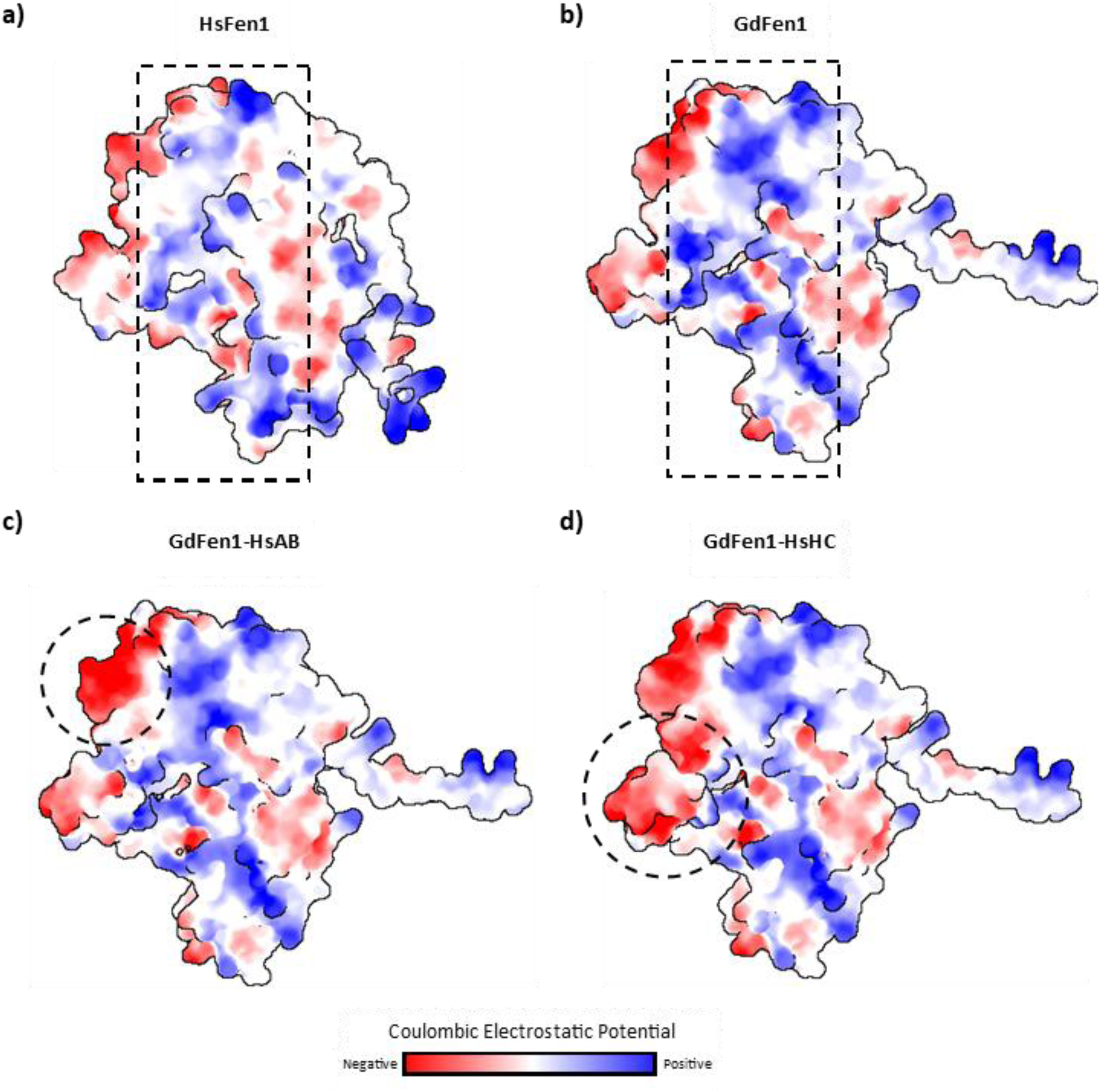
Electrostatic potential in GdFen1 mutants. The Coulombic Electrostatic Potential (ESP) is shown in the four predicted models. a) and b) Wildtype HsFen1 and GdFen1, the rectangles with dotted lines indicate the region of interaction with DNA. The last 13 residues on the HsFen1 carboxyl end have been deleted from the image to improve visualization. c) GdFen1-HsAB, dotted circle denotes the acid block d) GdFen1-HsHC, the dotted circle indicates the helical cap. The residues with negative potential are depicted in red while those with positive potential are shown in blue.

## 4. Discussion

Giardia duodenalis has several interesting features that have positioned it as a relevant and peculiar model organism to study those cellular mechanisms that are deeply understood in other organisms, such as bacteria or humans, but show an unexpected flexibility in this parasite. In this regard, G. duodenalis has been described under different circumstances as a minimalist organism, since it possesses a minimal kinome (Manning et al., 2011), a reduced apparatus for transcription, RNA processing, and most metabolic pathways, a small number of genes with introns as well as smaller intergenic sequences, as stated earlier this reduction of complexity, is likely associated to adoption of a parasitic lifestyle (Cernikova et al., 2018; Morrison et al., 2007). A similar pattern has been observed for mechanisms related to the maintenance of genomic stability, which is particularly remarkable since G. duodenalis exhibits a changing ploidy during its cell cycle from 4n to 16n (Bernander et al., 2001; Reiner et al., 2008), implying a higher susceptibility to DNA damage. In this context, an apparent absence of the error-prone non-homologous end joining (NHEJ) has been observed, since none of its elements have been detected with in silico approaches. The same is true for Nbs1, one of the three molecules constituting the MRN complex, a machinery involved in the first steps after the formation of a DSB (Reginato & Cejka, 2020). Nbs1 is essential for the functioning of the complex, however, though the participation of a giardial Mre11/Rad50 complex for in vivo homologous recombination in the parasite was confirmed, no Nbs1 was identified (Martinez-Miguel et al., 2017; Sandoval-Cabrera et al., 2015). Another interesting feature of G. duodenalis is the absence of a rad51 homologue, the main recombinase involved in the high-fidelity process pathway of homologous recombination repair. In contrast, G. duodenalis has two homologues of the DMC1 gene (DMC1A and DMC1B), which is involved in meiotic recombination in sexual organisms. It is noteworthy that this parasite is asexual, and no evidence of meiotic processes has yet been observed, however, DMC1B has been biochemically characterized and shown that upon DNA damage infliction it participates in the DNA repair process in the trophozoite (Torres-Huerta et al., 2016).Consistently, homologous recombination and single-strand annealing have also been shown to take place in G. duodenalis (Garcia-Lepe et al., 2022). All these interesting observations undoubtedly place this organism as a model for the study of DNA damage/repair.

Following this line of peculiar molecular features regarding genomic stability, this research aimed at unexplored pathways, with a focus on the giardial protein Fen1 (GdFen1). Using bioinformatics approaches, a feasible homologue for this gene was found in the Giardia genome, the resulting amino acid sequence showed conservation of the major domains: XPGN, XPGI and H2TH, involved in DNA binding and processing (Figure 1). These observations, along with a very similar spatial configuration of the protein compared to the human homologue, strongly implied that the GdFen1 might perform the biochemical activities described in other organisms, including humans (reviewed in (Balakrishnan & Bambara, 2013)). To test this hypothesis, two of the major functions of Fen1 were evaluated on the recombinant protein. As shown above, the parasite enzyme is able to bind and specifically cleave DNA structures bearing a 5’ flap (Figure 4a and c). Notably, this is in good agreement with previous reports demonstrating that Fen1 is able to trim single or double flap structures of different lengths in a sequence-independent manner (Balakrishnan & Bambara, 2013; Hohl et al., 2007; Tsutakawa et al., 2011). These findings confirm that G. duodenalis possesses a functional Fen1 nuclease that may be involved in DNA replication and repair in vivo, and also indirectly suggests that the BER pathway may be conserved in the parasite.

In vitro, Fen1 is able to bind and cleave various DNA structures, including double flaps, DNA nicking without a 5’ flap and also the processing of structures where the 5’ folds back to form a hairpin (Finger & Shen, 2010; Lyamichev et al., 1999). Indeed, it was previously reported that human Fen1 was able to bind but not cleave bubble structures (Hohl et al., 2007), as was corroborated in the EMSA performed with GdFen1, where it was capable of binding to the Bubble probe (Figure 5a). For these reasons, one of the most remarkable observations in this work was the unexpected ability of GdFen1 to cleave bubble structures (Figure 5b). When the Bubble probe was used, a similar result to that of the Flap probe was obtained, producing a DNA fragment bearing the fluorescent tag, but with a different size (Figure 3b). To confirm this observation and to exclude the possibility that the probes do not form the expected bubble structure, the same experiment was repeated with a completely different fluorescent bubble probe, where the same result was obtained (data not shown). It can be seen in Table 1 and Figure 3, that the same fluorescent oligonucleotide (A-bubble 5’ FAM*) was used to assemble both the Flap and Bubble probes, consequently, when the Flap probe is processed a detectable product of 20 nt must be produced (Figure 3a), however when the Bubble probe is cleaved it must generate a product of 40 nt long, including a hybrid of ssDNA (20 nt long) and dsDNA (20 nt long) (Figure 3b). This cleavage pattern was corroborated in all the nuclease experiments, where the Flap or Bubble probes clearly produce fragments of different sizes that fit the expected pattern. It is important to clarify that the use of bubble probes to study the cutting capacity of proteins within the Fen1 family in vitro does not require prior cleavage at the 5’-end of the bubble, as the XPG protein does when acting in the NER pathway in the cellular context, where XPF is required to perform a cleavage first to facilitate the second one by XPG (González-Corrochano et al., 2020; Hohl et al., 2007; Staresincic et al., 2009; Tsutakawa et al., 2011, 2020). These results provide compelling evidence for the special behavior of GdFen1 over bubble structures.

The ability to hydrolyze bubble structures has been widely attributed to the enzyme XPG, which has the same nuclease domains as Fen1, XPGN and XPGI (Scharer, 2013; Tsutakawa et al., 2020). However, the difference lies in the sequence between these two elements: while Fen1 contains a small sequence of about 70 amino acids, XPG has a considerably longer spacer region (R-domain) of about 680 amino acids. It has been conceived that the presence of this spacer is directly related to the ability of XPG to process bubble structures, as these are structurally larger (Hohl et al., 2007). Nonetheless, when this spacer segment was artificially inserted between the two nuclease domains in Fen1, the resulting Fen1-XPG hybrid protein did not fully replicate the ability of XPG to cleave bubble structures (Hohl et al., 2007). On the other hand, when the spacer region was eliminated from the XPG and replaced with the spacer from Fen1, XPG lost the capability of cleaving bubble structures (González-Corrochano et al., 2020; Tsutakawa et al., 2020). These contradictory observations indicate that although the presence of the spacer sequence is important for the biochemical activity of the nuclease, it does not fully explain its necessity for processing these DNA configurations, as was corroborated with GdFen1 which owns a short spacer region not dissimilar to other orthologues (Figure 1a and b).

In a separate study, crystallographic analyses were performed to elucidate the molecular basis of human Fen1 activities and to understand the substrate specificities exhibited by the 5’- nuclease superfamily (Tsutakawa et al., 2011). This research identified a specific structural loop, the “acid block,” a four-residue sequence of EEGE amino acids that is unique to Fen1 and absent from the other members of the superfamily. The acid block was predicted to act as an obstacle that prevents the processing of bubble structures by preventing DNA from passing beyond the pocket of the protein. Similarly, a second element was identified as critical for substrate cleavage, the “helical cap”, which assists in 5’-end selection and structurally forms the helical gateway, the entrance to the active site. Interestingly, when this helical cap from Fen1 was inserted into XPG and the original spacer region was replaced, it was observed that the processing capacity of the hybrid XPG on the bubble was reduced but not completely lost (Tsutakawa et al., 2011). The above results suggest two scenarios: first, the helical cap is partially involved in Fen1 specificity, and second, the spacer region of XPG is important but not fundamental for bubble DNA processing, similar to what was observed in the previous Fen1- XPG hybrid (Hohl et al., 2007). Regarding the results obtained in this study, as was observed in the sequence alignment (Figure 1a), none of the members of the Fornicata phylum included in the analysis have a full acid block sequence, some of them contain only one or two of the canonical amino acids of human Fen1 (Figure 1a, solid square). With respect to the helical cap, GdFen1 has a sequence that differs considerably from that of the human (Figure 1a, dotted square), specifically in the second helix of the helical cap (α5), variances that also confer a more positive charge potential to the local structure (Figure 7b). To investigate whether the differences in the acid block and helical cap are responsible for the cleavage activity performed on bubbles structures by GdFen1, two mutants were generated, where these sequences were replaced with the human counterpart. Interestingly, when the human acid block was inserted into the GdFen1, no affectations were detected in DNA binding or nuclease activity over flap or bubbles structures (Figure 5a, e and g), indicating that, in this particular case, the acid block is not acting as a constraint for bubble cleavage, in contrast to what was previously speculated (Tsutakawa et al., 2011). In agreement with these observations, the acid block mutation does not modify other features such as the gateway cavity or the electrostatic potential in a way that can impact the protein activities (Figure 6c and 7c). On the contrary, when the helical cap was mutated, GdFen1 lost nuclease capability, strongly suggesting that the giardial helical cap is directly involved in the cleavage of bubble structures (Figure 5d, e and g). Of note, this mutation additionally reduced the volume and shape of the helical gateway cavity, making it more like the human protein cavity and also creates a more negative surface in the helical cap, possibly impeding the access of the DNA to the active site (Figure 6d and 7d). This observation may also explain why the protein retained DNA binding activity, since any of these changes affect the main regions associated with DNA backbone interaction (González-Corrochano et al., 2020).

Considering all the peculiarities of G. duodenalis in terms of DNA repair and genomic stability, it is not entirely surprising that GdFen1 has an increased ability to recognize and process different substrates, which also fits the genetic minimalism of this parasite. The observations of this work not only confirm the functional capacity of G. duodenalis to process flap structures, relevant for the BER repair pathway and during the replicative process, but also suggest the presence of the NER pathway, whose existence had been ruled out in the parasite (Einarsson et al., 2015). In addition, these findings may also be an indication of some kind of redundancy between Fen1 and XPG. The ability to resolve voluptuous structures, such as bubbles, through the NER pathway becomes relevant in the parasite, as the use of UV radiation, which produces such lesions, is a widely used method for the removal of trophozoites and cysts for different purposes (Adeyemo et al., 2019; Li et al., 2008; Linden et al., 2002), although trophozoites are not commonly exposed to UV light due to the parasitic life cycle of the parasite (Bernander et al., 2001; Thompson et al., 1993). In addition to GdFen1, the presence of potential homologues of the NER pathway, including XPG, has been detected by our working group through bioinformatics approaches in G duodenalis. In vitro and in vivo characterization of these elements will help to improve the understanding of this pathway and how G. duodenalis resolves different types of DNA damage.

Despite the wealth of functional and structural studies on Fen1 and XPG, the helical cap remains an intriguing structure. So far, approaches based on hybrids of Fen1 with the spacer region of XPG or deletion of the same region in XPG have not been able to fully explain its role in substrate specificity. Furthermore, the structural disorganisation of the spacer region does not allow crystallographic studies to be performed on the full version of the XPG protein, which is a disadvantage for understanding the processing of the bubble structure. In conclusion, this work has investigated an unexplored element involved in genome maintenance and DNA repair in G. duodenalis. The results obtained provide evidence for the involvement of GdFen1 in the processing of flap structures, suggesting that the mechanisms by which this enzyme is used are conserved in the parasite. Despite the fact that Fen1 has been described as being unable to process bubble structures, GdFen1 does bind and cleave this type of DNA configuration, an ability that is possible due to variations in the amino acid sequence in the helical cap. These observations add to our knowledge of the resourcefulness of this organism in terms of genetic stability.

## Acknowledgements

We acknowledge to the Secretary of Education, Science, Technology and Innovation of the City of Mexico (SECTEI) for its contribution with the scholarship N° SECTEI/131/2023 to the postdoctoral researcher Ulises Omar García Lepe.

## 7. Author statement

Ulises Omar García-Lepe: Conceptualization, Formal Analysis, Investigation, Writing – Original Draft. Sofía Gabriela Tomás-Morales: Formal Analysis, Investigation. María Teresa Izaguirre- Hernández: Conceptualization, Formal Analysis, Investigation. María Luisa Bazán-Tejeda: Investigation, Supervision. Rosa María Bermúdez-Cruz: Conceptualization, Writing – Original Draft, Project administration, Funding Acquisition.

## References

Adeyemo, F. E., Singh, G., Reddy, P., Bux, F., & Stenstrom, T. A. (2019). Efficiency of chlorine and UV in the inactivation of Cryptosporidium and Giardia in wastewater. PLoS One, 14(5), e0216040. 10.1371/journal.pone.0216040

Balakrishnan, L., & Bambara, R. A. (2013). Flap endonuclease 1. Annu Rev Biochem, 82, 119–138. 10.1146/annurev-biochem-072511-122603

Bernander, R., Palm, J. E., & Svard, S. G. (2001). Genome ploidy in different stages of the Giardia lamblia life cycle. Cell Microbiol, 3(1), 55–62. 10.1046/j.1462-5822.2001.00094.x

Cernikova, L., Faso, C., & Hehl, A. B. (2018). Five facts about Giardia lamblia. PLoS Pathog, 14(9), e1007250. 10.1371/journal.ppat.1007250

Dehé, P. M., & Gaillard, P. H. L. (2017). Control of structure-specific endonucleases to maintain genome stability. Nature Reviews. Molecular Cell Biology, 18(5), 315–330. 10.1038/NRM.2016.177

Dehe, P. M., & Gaillard, P. H. L. (2017). Control of structure-specific endonucleases to maintain genome stability. Nat Rev Mol Cell Biol, 18(5), 315–330. 10.1038/nrm.2016.177

Einarsson, E., Svard, S. G., & Troell, K. (2015). UV irradiation responses in Giardia intestinalis. Exp Parasitol, 154, 25–32. 10.1016/j.exppara.2015.03.024

Fagbemi, A. F., Orelli, B., & Scharer, O. D. (2011). Regulation of endonuclease activity in human nucleotide excision repair. DNA Repair (Amst), 10(7), 722–729. 10.1016/j.dnarep.2011.04.022

Finger, L. D., & Shen, B. %J A. G. C. O. H. (2010). FEN1 (flap structure-specific endonuclease 1).

Garcia-Lepe, U. O., Espinoza-Corona, S., Bazan-Tejeda, M. L., Nunez-Jurado, F. M., & Bermudez-Cruz, R. M. (2022). Giardia duodenalis carries out canonical homologous recombination and single-strand annealing. Res Microbiol, 173(8), 103984. 10.1016/j.resmic.2022.103984

González-Corrochano, R., Ruiz, F. M., Taylor, N. M. I., Huecas, S., Drakulic, S., Spínola-Amilibia, M., & Fernández-Tornero, C. (2020). The crystal structure of human XPG, the xeroderma pigmentosum group G endonuclease, provides insight into nucleotide excision DNA repair. Nucleic Acids Research, 48(17), 9943–9958. 10.1093/NAR/GKAA688

Gouet, P., Robert, X., & Courcelle, E. (2003). ESPript/ENDscript: Extracting and rendering sequence and 3D information from atomic structures of proteins. Nucleic Acids Res, 31(13), 3320–3323. 10.1093/nar/gkg556

Hohl, M., Dunand-Sauthier, I., Staresincic, L., Jaquier-Gubler, P., Thorel, F., Modesti, M., Clarkson, S. G., & Scharer, O. D. (2007). Domain swapping between FEN-1 and XPG defines regions in XPG that mediate nucleotide excision repair activity and substrate specificity. Nucleic Acids Res, 35(9), 3053–3063. 10.1093/nar/gkm092

Jumper, J., Evans, R., Pritzel, A., Green, T., Figurnov, M., Ronneberger, O., Tunyasuvunakool, K., Bates, R., Žídek, A., Potapenko, A., Bridgland, A., Meyer, C., Kohl, S. A. A., Ballard, A. J., Cowie, A., Romera-Paredes, B., Nikolov, S., Jain, R., Adler, J., … Hassabis, D. (2021). Highly accurate protein structure prediction with AlphaFold. Nature 2021 596:7873, 596(7873), 583–589. 10.1038/s41586-021-03819-2

Katoh, K., Standley, D. M. %J M. biology, & evolution. (2013). MAFFT multiple sequence alignment software version 7: improvements in performance and usability. 30(4), 772–780.

Li, D., Craik, S. A., Smith, D. W., & Belosevic, M. (2008). Survival of Giardia lamblia trophozoites after exposure to UV light. FEMS Microbiol Lett, 278(1), 56–61. 10.1111/j.1574-6968.2007.00972.x

Lieber, M. R. (1997). The FEN-1 family of structure-specific nucleases in eukaryotic DNA replication, recombination and repair. Bioessays, 19(3), 233–240. 10.1002/bies.950190309

Linden, K. G., Shin, G. A., Faubert, G., Cairns, W., & Sobsey, M. D. (2002). UV disinfection of Giardia lamblia cysts in water. Environ Sci Technol, 36(11), 2519–2522. 10.1021/es0113403

Liu, Y., Kao, H. I., & Bambara, R. A. (2004). Flap endonuclease 1: a central component of DNA metabolism. Annu Rev Biochem, 73, 589–615. 10.1146/annurev.biochem.73.012803.092453

Lyamichev, V., Brow, M. A., Varvel, V. E., & Dahlberg, J. E. (1999). Comparison of the 5’ nuclease activities of taq DNA polymerase and its isolated nuclease domain. Proc Natl Acad Sci U S A, 96(11), 6143–6148. 10.1073/pnas.96.11.6143

Manning, G., Reiner, D. S., Lauwaet, T., Dacre, M., Smith, A., Zhai, Y., Svard, S., & Gillin, F. D. (2011). The minimal kinome of Giardia lamblia illuminates early kinase evolution and unique parasite biology. Genome Biol, 12(7), R66. 10.1186/gb-2011-12-7-r66

Marchler-Bauer, A., Derbyshire, M. K., Gonzales, N. R., Lu, S., Chitsaz, F., Geer, L. Y., Geer, R. C., He, J., Gwadz, M., Hurwitz, D. I., Lanczycki, C. J., Lu, F., Marchler, G. H., Song, J. S., Thanki, N., Wang, Z., Yamashita, R. A., Zhang, D., Zheng, C., & Bryant, S. H. (2015). CDD: NCBI’s conserved domain database. Nucleic Acids Res, 43(Database issue), D222-6. 10.1093/nar/gku1221

Martinez-Miguel, R. M., Sandoval-Cabrera, A., Bazan-Tejeda, M. L., Torres-Huerta, A. L., Martinez-Reyes, D. A., & Bermudez-Cruz, R. M. (2017). Giardia duodenalis Rad52 protein: biochemical characterization and response upon DNA damage. J Biochem, 162(2), 123–135. 10.1093/jb/mvx009

Meng, E. C., Goddard, T. D., Pettersen, E. F., Couch, G. S., Pearson, Z. J., Morris, J. H., & Ferrin, T. E. (2023). UCSF ChimeraX: Tools for structure building and analysis. Protein Science, 32(11), e4792. 10.1002/PRO.4792

Morrison, H. G., McArthur, A. G., Gillin, F. D., Aley, S. B., Adam, R. D., Olsen, G. J., Best, A. A., Cande, W. Z., Chen, F., Cipriano, M. J., Davids, B. J., Dawson, S. C., Elmendorf, H. G., Hehl, A. B., Holder, M. E., Huse, S. M., Kim, U. U., Lasek-Nesselquist, E., Manning, G., … Sogin, M. L. (2007). Genomic minimalism in the early diverging intestinal parasite Giardia lamblia. Science, 317(5846), 1921–1926. 10.1126/science.1143837

Ordoñez-Quiroz, A., Ortega-Pierres, M. G., Bazán-Tejeda, M. L., & Bermúdez-Cruz, R. M. (2018). DNA damage induced by metronidazole in Giardia duodenalis triggers a DNA homologous recombination response. Experimental Parasitology, 194, 24–31. 10.1016/J.EXPPARA.2018.09.004

Pettersen, E. F., Goddard, T. D., Huang, C. C., Meng, E. C., Couch, G. S., Croll, T. I., Morris, J. H., & Ferrin, T. E. (2021). UCSF ChimeraX: Structure visualization for researchers, educators, and developers. Protein Science : A Publication of the Protein Society, 30(1), 70–82. 10.1002/PRO.3943

Reginato, G., & Cejka, P. (2020). The MRE11 complex: A versatile toolkit for the repair of broken DNA. DNA Repair (Amst), 91–92, 102869. 10.1016/j.dnarep.2020.102869

Reiner, D. S., Ankarklev, J., Troell, K., Palm, D., Bernander, R., Gillin, F. D., Andersson, J. O., & Svard, S. G. (2008). Synchronisation of Giardia lamblia: identification of cell cycle stage-specific genes and a differentiation restriction point. Int J Parasitol, 38(8–9), 935– 944. 10.1016/j.ijpara.2007.12.005

Sandoval-Cabrera, A., Zarzosa-Alvarez, A. L., Martinez-Miguel, R. M., & Bermudez-Cruz, R. M. (2015). MR (Mre11-Rad50) complex in Giardia duodenalis: In vitro characterization and its response upon DNA damage. Biochimie, 111, 45–57. 10.1016/j.biochi.2015.01.008

Sayers, E. W., Beck, J., Bolton, E. E., Brister, J. R., Chan, J., Connor, R., Feldgarden, M., Fine, A. M., Funk, K., Hoffman, J., Kannan, S., Kelly, C., Klimke, W., Kim, S., Lathrop, S., Marchler-Bauer, A., Murphy, T. D., O’Sullivan, C., Schmieder, E., … Pruitt, K. D. (2024). Database resources of the National Center for Biotechnology Information in 2025. Nucleic Acids Research, 53(D1), D20. 10.1093/NAR/GKAE979

Scharer, O. D. (2013). Nucleotide excision repair in eukaryotes. Cold Spring Harb Perspect Biol, 5(10), a012609. 10.1101/cshperspect.a012609

Spivak, G. (2015). Nucleotide excision repair in humans. DNA Repair (Amst), 36, 13–18. 10.1016/j.dnarep.2015.09.003

Staresincic, L., Fagbemi, A. F., Enzlin, J. H., Gourdin, A. M., Wijgers, N., Dunand-Sauthier, I., Giglia-Mari, G., Clarkson, S. G., Vermeulen, W., & Schärer, O. D. (2009). Coordination of dual incision and repair synthesis in human nucleotide excision repair. The EMBO Journal, 28(8), 1111. 10.1038/EMBOJ.2009.49

Thompson, R. C. A., Reynoldson, J. A., & Mendis, A. H. W. (1993). Giardia and Giardiasis. 32, 71–160. 10.1016/S0065-308X(08)60207-9

Torres-Huerta, A. L., Martinez-Miguel, R. M., Bazan-Tejeda, M. L., & Bermudez-Cruz, R. M. (2016). Characterization of recombinase DMC1B and its functional role as Rad51 in DNA damage repair in Giardia duodenalis trophozoites. Biochimie, 127, 173–186. 10.1016/j.biochi.2016.05.014

Tsutakawa, S. E., Classen, S., Chapados, B. R., Arvai, A. S., Finger, L. D., Guenther, G., Tomlinson, C. G., Thompson, P., Sarker, A. H., Shen, B., Cooper, P. K., Grasby, J. A., & Tainer, J. A. (2011). Human flap endonuclease structures, DNA double-base flipping, and a unified understanding of the FEN1 superfamily. Cell, 145(2), 198–211. 10.1016/j.cell.2011.03.004

Tsutakawa, S. E., Sarker, A. H., Ng, C., Arvai, A. S., Shin, D. S., Shih, B., Jiang, S., Thwin, A. C., Tsai, M. S., Willcox, A., Her, M. Z., Trego, K. S., Raetz, A. G., Rosenberg, D., Bacolla, A., Hammel, M., Griffith, J. D., Cooper, P. K., & Tainer, J. A. (2020). Human XPG nuclease structure, assembly, and activities with insights for neurodegeneration and cancer from pathogenic mutations. Proc Natl Acad Sci U S A, 117(25), 14127–14138. 10.1073/pnas.1921311117

Zhang, Y., & Skolnick, J. (2005). TM-align: a protein structure alignment algorithm based on the TM-score. Nucleic Acids Research, 33(7), 2302–2309. 10.1093/NAR/GKI524

